## Supplementary Figures for "Differences in phenotype between long-lived memory B cells against *Plasmodium falciparum* merozoite antigens and variant surface antigens"

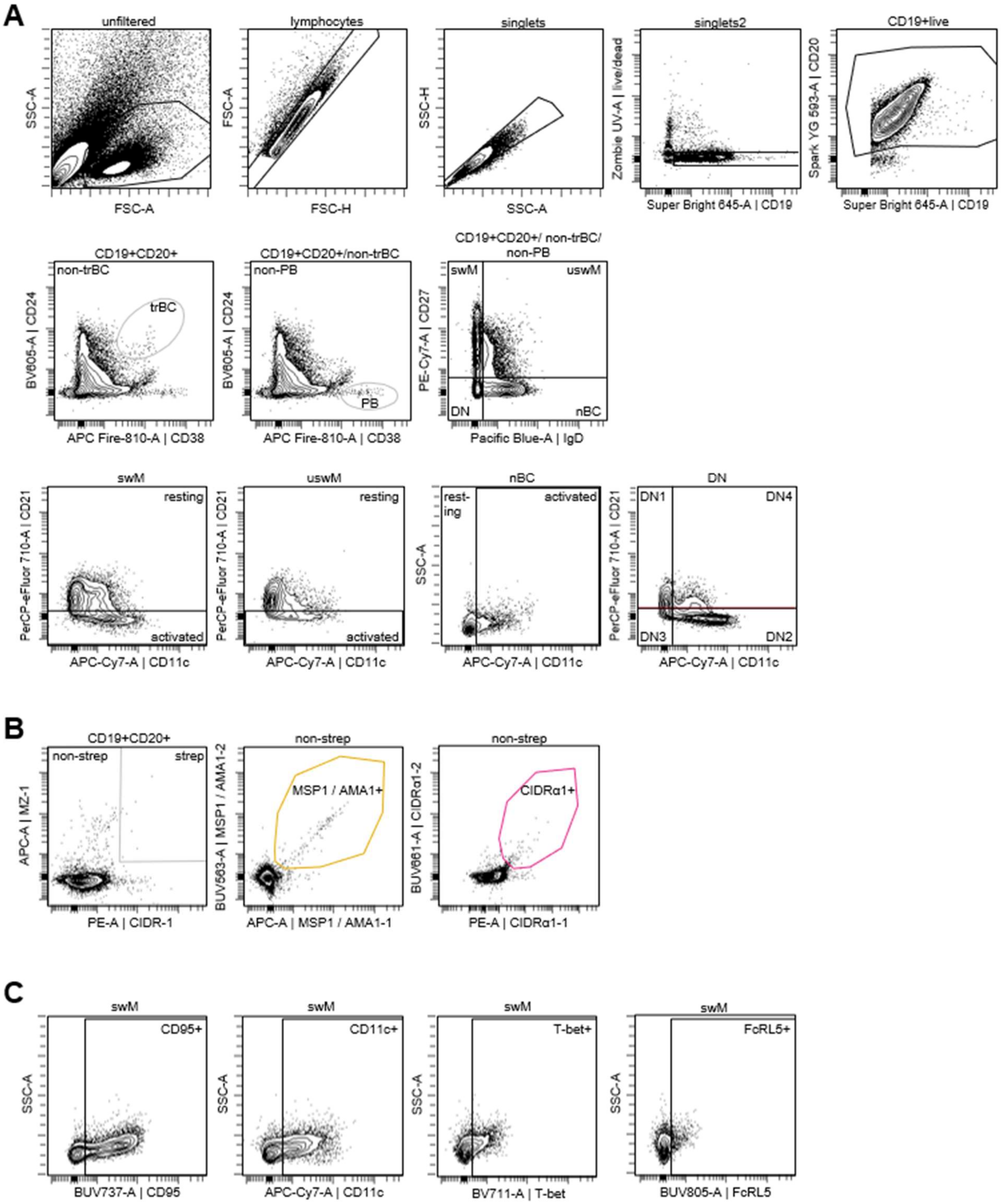


**Figure S1: Gating strategy. A)** The gating scheme used to identify resting and activated B cell populations, as well as subpopulations of double negative cells. **B)** The gating strategy used to identify antigen-specific B cells. **C)** The gating strategy used to identify CD95$^+$, CD11c$^+$, T-bet$^+$, and FcRL5$^+$ B cells. trBC, transitional B cells; PB, plasmablasts; swM, switched memory; unswM, unswitched memory; nBC, naïve B cells; DN, double negative.

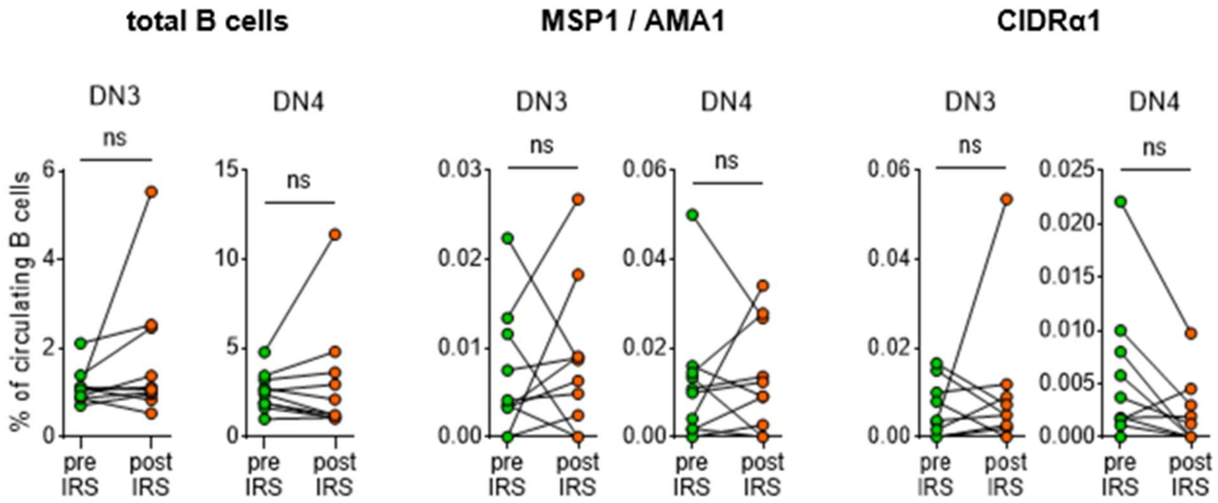


**Figure S2: The percentage of DN3 and DN4 B cells among circulating B cells.** The percentages are shown for total B cells (left), MSP1/AMA1-specific B cells (middle), and CIDRα1-specific B cells (right). Differences between groups were evaluated using a Wilcoxon matched-pairs signed-rank test. DN, double negative; ns, not significant.

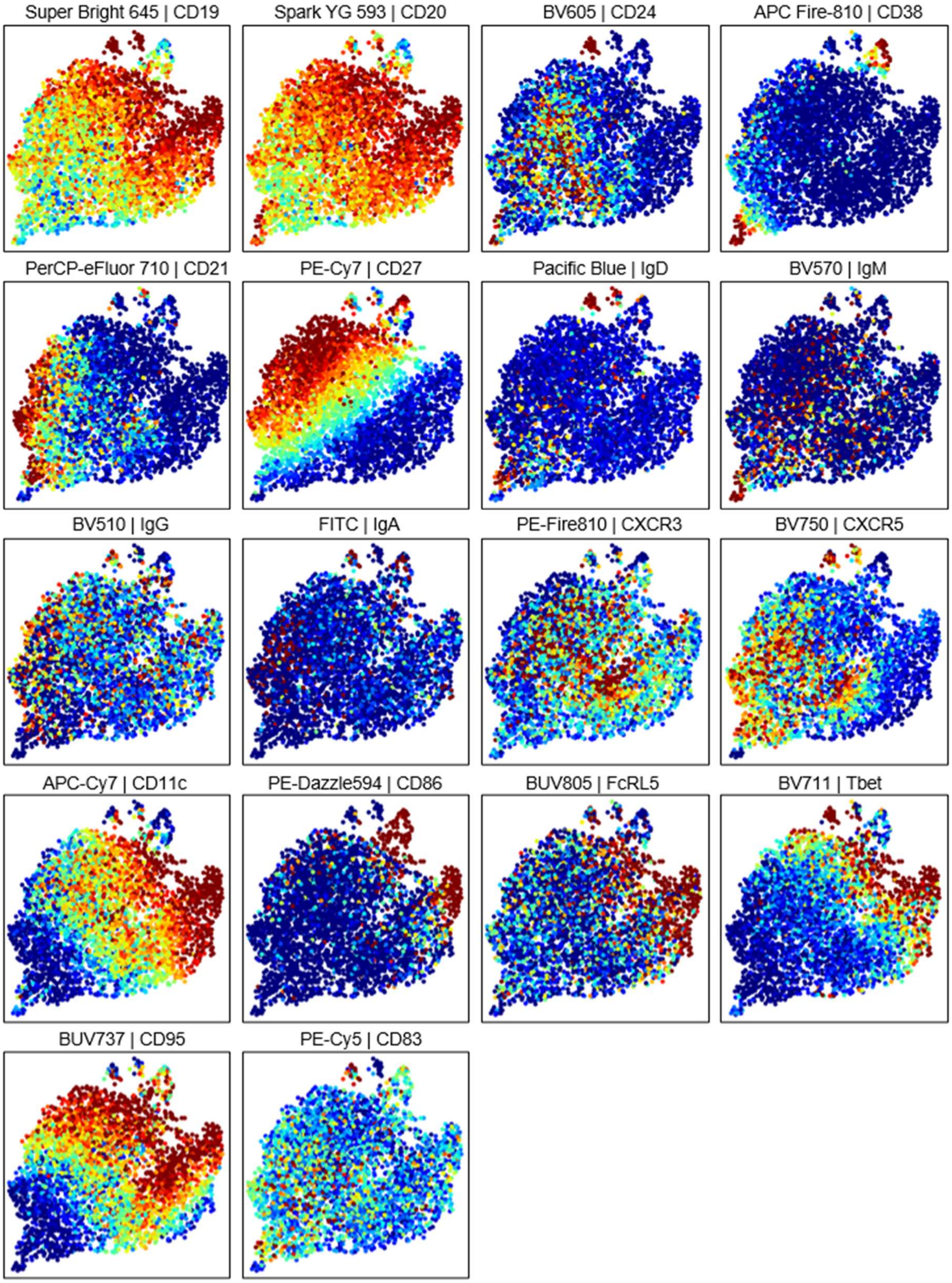


**Figure S3: Projection of the surface and intracellular markers used to generate the composite UMAP of antigen-specific B cells onto this UMAP.**

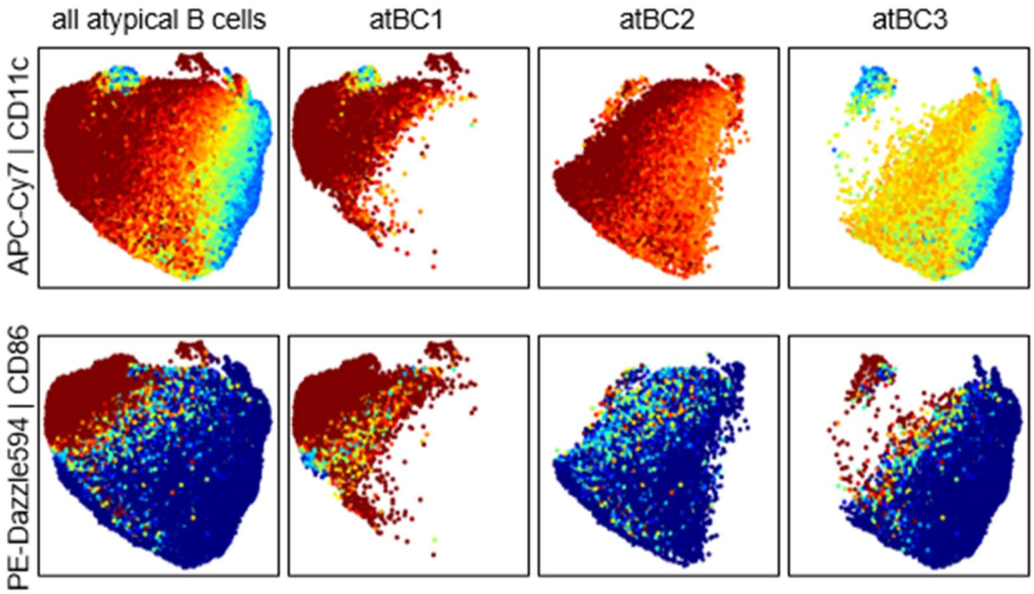


**Figure S4: CD11c and CD86 expression projected onto the UMAP of all atypical B cells as well as each of the three subsets individually.**

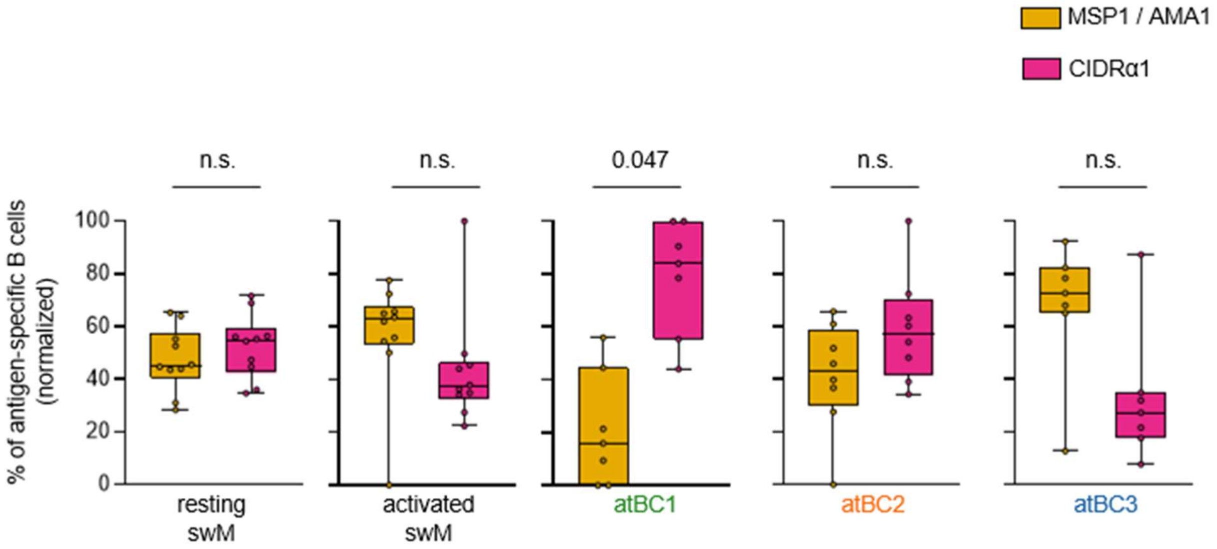


**Figure S5: The normalized percentage of MSP1/AMA1-specific and CIDRα1-specific B cells among different B cell subsets.** Values were calculated for individual donors. For the three atypical B cell subsets, donors were only included if the total size of the subset allowed for the detection of at least one MSP1/AMA1-specific B cell and one CIDRα1-specific B cell. For example, if the total percentage of MSP1/AMA1-specific B cells is 1.5%, a B cell population needs to contain at least 1 / 1.5 * 100 = 67 cells to be included in this analysis. Because atypical B cell subsets in samples obtained post-IRS were too small to perform this analysis, only pre-IRS data is shown. swM, switched memory B cell; atBC*i*, atypical B cell subset *i*. n.s., not significant.
